# SARS-CoV-2 Infection of Microglia Elicits Pro-inflammatory Activation and Apoptotic Cell Death

**DOI:** 10.1101/2022.01.04.475015

**Authors:** Gi Uk Jeong, Jaemyun Lyu, Kyun-Do Kim, Young Cheul Chung, Gun Young Yoon, Sumin Lee, Insu Hwang, Won-Ho Shin, Junsu Ko, June-Yong Lee, Young-Chan Kwon

## Abstract

Accumulating evidence suggests that severe acute respiratory syndrome coronavirus 2 (SARS-CoV-2) infection causes various neurological symptoms in coronavirus disease 2019 (COVID-19) patients. The most dominant immune cells in the brain are microglia. Yet, the relationship between neurological manifestations, neuroinflammation, and host immune response of microglia to SARS-CoV-2 has not been well characterized. Here, we report that SARS-CoV-2 can directly infect human microglia, eliciting M1-like pro-inflammatory responses, followed by cytopathic effects. Specifically, SARS-CoV-2 infected human microglial clone 3 (HMC3), leading to inflammatory activation and cell death. RNA-seq analysis also revealed that ER stress and immune responses were induced in the early and apoptotic processes in the late phase of viral infection. SARS-CoV-2-infected HMC3 showed the M1 phenotype and produced pro-inflammatory cytokines such as interleukin (IL)-1β, IL-6, and tumour necrosis factor α (TNF-α), but not the anti-inflammatory cytokine IL-10. After this pro-inflammatory activation, SARS-CoV-2 infection promoted both intrinsic and extrinsic death receptor-mediated apoptosis in HMC3. Using K18-hACE2 transgenic mice, murine microglia were also infected by intranasal inoculation of SARS-CoV-2. This infection induced the acute production of pro-inflammatory microglial IL-6 and TNF-α and provoked a chronic loss of microglia. Our findings suggest that microglia are potential mediators of SARS-CoV-2-induced neurological problems and, consequently, can be targets of therapeutic strategies against neurological diseases in COVID-19 patients.

**IMPORTANCE:** Recent studies reported neurological manifestations and complications in COVID-19 patients, which are associated with neuroinflammation. As microglia are the dominant immune cells in brains, it needs to be elucidate the relationship between neuroinflammation and host immune response of microglia to SARS-CoV-2. Here, we suggest that SARS-CoV-2 can directly infect human microglia with cytopathic effect (CPE) using human microglial clone 3 (HMC3). The infected microglia were promoted to pro-inflammatory activation following apoptotic cell death. This pro-inflammatory activation was accompanied by the high production of pro-inflammatory cytokines, and led to neurotoxic-M1 phenotype polarization. *In vivo*, murine microglia were infected and produced pro-inflammatory cytokines and provoked a chronic loss using K18-hACE2 mice. Thus, our data present that SARS-CoV-2-infected microglia are potential mediators of neurological problems in COVID-19 patients. In addition, HMC3 cells are susceptible to SARS-CoV-2 and exhibit the CPE, which can be further used to investigate cellular and molecular mechanisms of neuroinflammation reported in COVID-19 patients.

## INTRODUCTION

Coronaviruses, which are enveloped positive-single-stranded RNA viruses that belong to the *Coronaviridae* family, cause mild to severe respiratory, enteric, and neurological diseases in humans and animals (1). At the end of 2019, a pneumonia outbreak caused by severe acute respiratory syndrome coronavirus 2 (SARS-CoV-2) was reported in Wuhan, China, and this novel coronavirus gave rise to the current coronavirus disease 2019 (COVID-19) pandemic (2). By cross-species transmission to human, as of December 2021, regarding the COVID-19 situation reports from WHO, globally over 280 million COVID-19 confirmed cases and 5 million deaths have been reported.

Although COVID-19 is primarily characterized as a respiratory disease, multiple organ dysfunction syndromes may occur in several organs, including the brain, which contribute to neurological manifestations (3). Acute neurological and psychiatric complications of COVID-19 often occurred even in persons younger than 60 years of age (4). Moreover, cortical signal alteration (5), loss of white matter, and axonal injury (6) have been reported, as well as increasing observations of neurological issues, including headache, ischemic stroke, seizures, delirium, anosmia, ageusia, encephalopathy, and total paralysis (7-14). Patients with more severe infection are more likely to have neurological manifestations and impairment and are at a higher risk of mortality (15).

Microglia are macrophage-like brain’s immune cells in the central nervous system (CNS). They have key functions in maintaining brain homeostasis and in the rapid response to injury and inflammation (16). When microglia respond to immunological stimuli, they become activated and transform from a ramified into an amoeboid morphology, releasing interleukin (IL)-1β, IL-6, and tumour necrosis α (TNF-α) (17). Activated microglia consist of a dual phenotype, wherein M1 or the classically activated state is neurotoxic and involved in neuroinflammation, and M2 or the alternatively activated state is neuroprotective (18-20). Increasing evidence suggests that the over-activation and dysregulation of microglia might result in disastrous and progressive neurotoxic consequences (21-24). In brains of deceased COVID-19 patients, microgliosis and immune cell accumulation were observed (25), as well as microglial nodules caused by massive microglia activation in the medulla oblongata (26) and cerebellar dentate nuclei (27). The neuroinvasive capacity (28) and olfactory transmucosal invasion of SARS-CoV-2 in COVID-19 patients (29) have also been reported. Additionally, human microglia express SARS-CoV-2 entry factors such as angiotensin-converting enzyme 2 (ACE2) and transmembrane protease serine subtype 2 (TMPRSS2) (30). Thus, we hypothesised that microglial activation by direct SARS-CoV-2 infection could be one of the major mechanisms driving the neuroinflammation and neurological complications.

Despite accumulating evidence, little is known regarding the mechanisms involved in the neuroinflammation of SARS-CoV-2 infection. In this study, we demonstrated that SARS-CoV-2 can directly infect human microglia and induce pro-inflammatory responses, reflecting polarization toward the M1 phenotype. We further showed that the SARS-CoV-2 infection led to apoptosis as the cytopathic effect (CPE) through both intrinsic and extrinsic apoptotic pathways. Moreover, we found that murine microglia expressing hACE2 were infected by intranasally administered SARS-CoV-2, followed by microglial pro-inflammatory activation and loss of their population.

## RESULTS

### SARS-CoV-2 directly infects a human microglial cell line with cytopathic effects

In a post-mortem study, more than half of 25 COVID-19 patients presented high microglial immune activation with microglial nodules, and immune filtration was linked to axonal damage (31). Another post-mortem study reported that viral nucleocapsid protein (NP) was localized into microglia of the brain cortex in deceased COVID-19 patients (32). To investigate whether SARS-CoV-2 can infect human microglia, we used immortalized human embryonic brain-derived primary microglia (HMC3). This cell line was infected with one multiplicity of infection (MOI) of SARS-CoV-2, along with other well-known SARS-CoV-2 susceptible cell lines, including Caco-2, which is derived from human colon carcinoma, and Vero E6, which is a kidney epithelial cell line from an African green monkey (33). Viral RNA was detected by quantitative real-time polymerase chain reaction (RT-qPCR) and was slightly increased in HMC3 in a time-dependent manner, despite the lower viral copies per microgram of the total RNA in HMC3 than that of Caco-2 and Vero E6 (Fig. 1A). When secreted virus particles were measured by focus forming assay, the progeny viruses were detected in line with intracellular viral RNA at a much lower level than that of Caco-2 and Vero E6 (Fig. 1B). Notably, the intracellular viral RNA at 2 dpi was reduced by CR3022 neutralizing antibody in a dose-dependent manner (Fig. 1C). To confirm that SARS-CoV-2 infects HMC3, the three cell lines were infected as before, followed by immunofluorescence assay to detect dsRNA and NP at 3 dpi, verified by an additional detection of S and NP (Fig. 1D, Supplementary Fig. 1). Also, the cell lysates at 3 dpi were analysed by western blotting to detect NP (Fig. 1E). Importantly, SARS-CoV-2 infection of HMC3 triggered cell death as one of the CPE (Fig. 1F), similar to that in Vero E6, which has been known to exhibit the CPE, but not Caco-2 (33) (Supplementary Fig. 2). Beginning at 4 dpi, infected HMC3 led to cell death, and by 6 dpi, most cells had died. Crystal violet stating at 4 and 6 dpi showed that the cell-covered areas were diminished but those of 4 and 6 dpi were significantly recovered by CR3022 neutralizing antibody (Fig. 1G). Thus, these findings proposed that HMC3 is another human cell line, which is susceptible to SARS-CoV-2 infection and exhibits the CPE.

**Figure 1.**
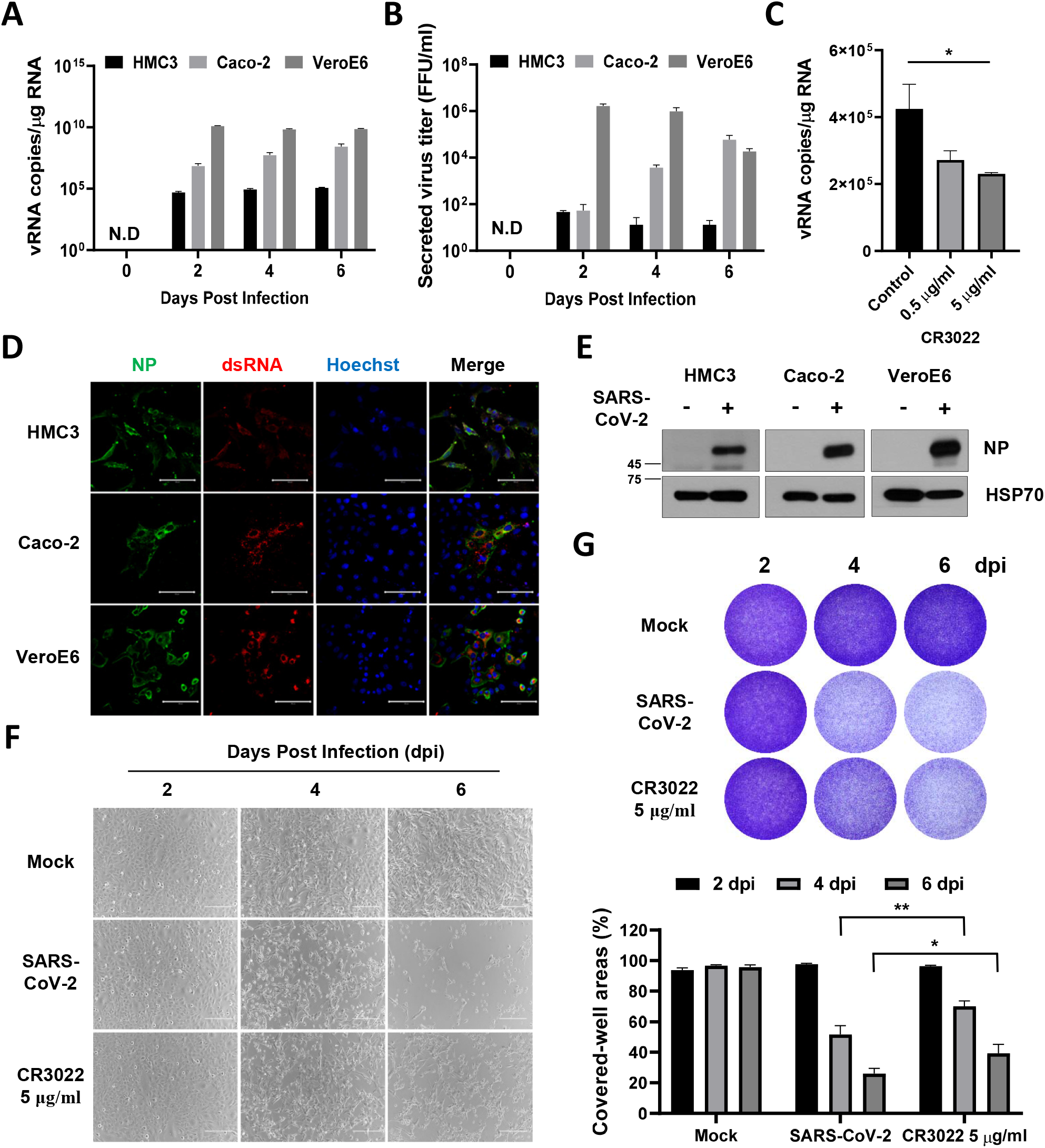
SARS-CoV-2 directly infects human microglia cells, eliciting CPE. **A** HMC3, Caco-2, and Vero E6 cells were infected with one MOI of SARS-CoV-2. The total cellular RNA was extracted at 2, 4, and 6 dpi to detect the viral RNA of the SARS-CoV-2 NP gene by Quantitative real-time polymerase chain reaction (RT-qPCR). The graph shows viral RNA copies per microgram of total cellular RNA on each day. **B** The culture media derived from SARS-CoV-2-infected cells were serially diluted and used for focus forming assay. The graph shows the secreted virus titre as focus forming units (FFU). **C** The graph shows viral RNA copies per microgram of total cellular RNA at 2 dpi after treatment with the increasing amount of CR3022 neutralizing antibody. **D** Confocal images of SARS-CoV-2-infected HMC3 (top row), Caco-2 (middle row), and Vero E6 (bottom row), demonstrating infection of these cells by immunofluorescence assay with anti-SARS-CoV-2 NP and anti-dsRNA antibodies. Scale bar = 100 μm. **E** Western blotting of SARS-CoV-2 NP in each infected cell. The 70-KDa heat shock protein (Hsp70) served as the loading control. **F** Phase-contrast images of the mock or SARS-CoV-2-infected HMC3 in the absence/presence of CR3022 neutralizing antibody at 2, 4, and 6 dpi, indicating cell death as the CPE by microscopy. Scale bar = 200 μm. **G** Images of crystal violet staining of the mock or SARS-CoV-2-infected HMC3 in the absence/presence of CR3022 neutralizing antibody, plated in the 12-well (upper). The graph shows the percent measurements of crystal violet-stained cell covered areas by ImmunoSpot reader (lower). Statistically significant differences between the groups were determined by Student’s t-test; \**P* < 0.05; \*\**P* < 0.01. Symbols represent mean ± SEM.

### Distinct transcriptional signatures and gene expression changes in SARS-CoV-2-infected HMC3

To assess the effect of SARS-CoV-2 infection on gene expression in HMC3, we performed RNA sequencing (RNA-seq) analysis on SARS-CoV-2-infected HMC3 at 3 and 6 dpi (S3 and S6, respectively), and an uninfected control (Mock, M), for comparison. Overall, the gene expression data indicated that M and S3 had relatively similar transcriptional signatures, while S6 had different signatures both in fold-change, and adjusted P-value (Fig. 2A, 2B). Differentially expressed gene (DEG) counts largely increased from 67 (S3) to 2,342 (S6) (Fig. 2C). More than half of the S3 DEGs overlapped (52.2 %, 35/67 DEGs) with S6, and most these genes were upregulated both at S3 and S6 (88.57 %, 31/35 overlapped DEGs) (Supplementary Fig. 3A). These results suggest that gene expression changes induced by SARS-CoV-2-infection increased with time, and accelerated more from S3 to S6, than from M to S3.

**Figure 2.**
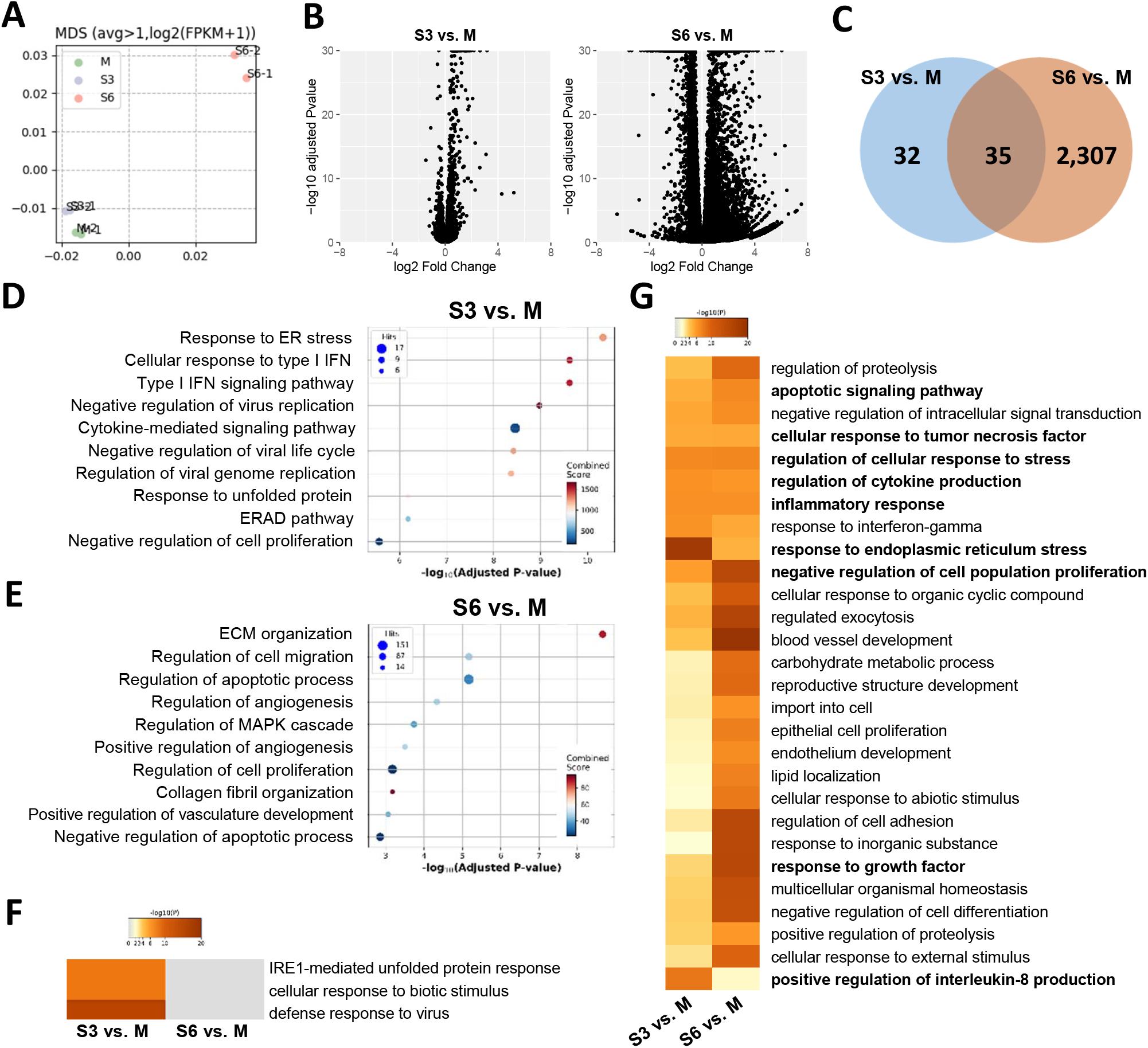
RNA-sequencing analysis of SARS-CoV-2-infected HMC3 cells. **A** Multidimensional analysis of genes expressed over one mean fragment per kilobase per million mapped fragments (FPKM). **B** Volcano plots of SARS-CoV-2-infected HMC3 cells at 3 dpi (S3; left) and 6 dpi (S6; right) compared to mock (M). **C** Venn diagram of differently expressed genes (DEGs) at S3 and S6. **D-E** Top 10 gene ontology (GO) enrichment terms for (D) S3 and (E) S6 DEGs. **F-G** Heatmaps showing GO enrichment terms for S3 and S6, significantly changed in both (F) S3 and (G) S6.

According to the over-representation analysis (ORA) of the gene ontology (GO) biological process, S3 DEGs in the early phase of viral infection were highly enriched for endoplasmic reticulum (ER) stress conditions, and antiviral immune responses, including type I interferons (IFNs) and cytokine-mediated signaling pathways (Fig. 2D). Given that RNA viruses induce ER stress through viral polypeptides and immune responses by double-stranded-RNA intermediates (34, 35), this result would imply SARS-CoV-2 infection and replication in HMC3. On the other hand, S6 DEGs were highly enriched for apoptotic processes (Fig. 2E). Although ER stress-related terms were not listed up in Fig. 2E, most of upregulated DEGs both in S3 and S6 (DNAJB7, DDIT3, HSPA5, HSP90B1, HYOU1, PDIA4, SEL1L) were highly enriched for ER stress, indicating that ER stress is still induced at S6 (Supplementary Fig. 3B). Given that prolonged ER stress ensues apoptosis (36), these results imply that defense responses against SARS-CoV-2 infection are elicited in the early phase of the viral infection, later gene expression changes contribute to apoptosis. In line with this, heatmap and network analyses of GO biological processes using Metascape (Fig. 2F, 2G, Supplementary Fig. 4), immune responses, cytokine production, ER stress responses, and defense responses to virus were distinctly enriched in S3, while apoptotic signaling pathway, negative regulation of cell population, growth factor response were significantly altered in S6. To conclude, SARS-CoV-2 infection in HMC3 induced gene expression of antiviral immune and ER stress responses in the early phase (S3) and of apoptosis in the late phase (S6).

### Microglial pro-inflammatory activation and M1 phenotype polarization by SARS-CoV-2-infection

In response to viral infection, microglia are activated and polarized into the pro-inflammatory M1 phenotype or the anti-inflammatory M2 phenotype (37). To assess the effects of SARS-CoV-2 infection on microglial activation and polarization in HMC3, we analysed the DEGs associated with immune response and microglial polarization. Several genes related to type I IFNs and innate immunity such as MXs, the 2’,5’-oligoadenylate synthetases (OASs), and Complement component 3 (C3) were upregulated at 3 or 6 dpi in the RNA-seq data (Fig. 3A). The M1 phenotype polarization related genes such as IL-1β, IL-6, and CXCL1 also showed increased RNA expression levels (Fig. 3B), indicating that SARS-CoV-2 evoked pro-inflammatory activation and polarization toward the M1 phenotype in HMC3. To confirm this, we assessed the expression of ILs and IFNs by RT-qPCR (Supplementary Fig. 5) and detected secreted cytokines in culture media by enzyme-linked immunosorbent assay (ELISA) (Fig. 3C). Pro-inflammatory cytokines such as IL-1β, IL-6, and TNF-α, but not the immune regulatory cytokine IL-10, were highly produced and secreted by SARS-CoV-2 infection. CD68 is a lysosomal protein expressed in high levels by macrophages and activated microglia and in low levels by resting microglia (38). The protein expression level of CD68 was increased in infected HMC3 (Fig. 3D), along with other makers of microglial pro-inflammatory activation such as CX3CL1 and CX3CR1 (Fig. 3E). These results evidently indicate that SARS-CoV-2-infected HMC3 are pro-inflammatory activated. To verify whether activated microglia polarized into the M1 phenotype, we estimated the RNA and protein expressions of the M1 phenotype markers. When we assessed the RNA expression level of each representative marker of the M1 (NOS2) and M2 phenotypes (Arigase-1), only NOS2 was highly expressed at 6 dpi (Fig. 3F). Protein expression levels of M1 markers, CD16, phospho-Stat1, and Stat1, were also induced at 4 and 6 dpi (Fig. 3G). Therefore, by SARS-CoV-2 infection, HMC3 was activated and polarized into the inflammatory M1 phenotype, producing pro-inflammatory cytokines.

**Figure 3.**
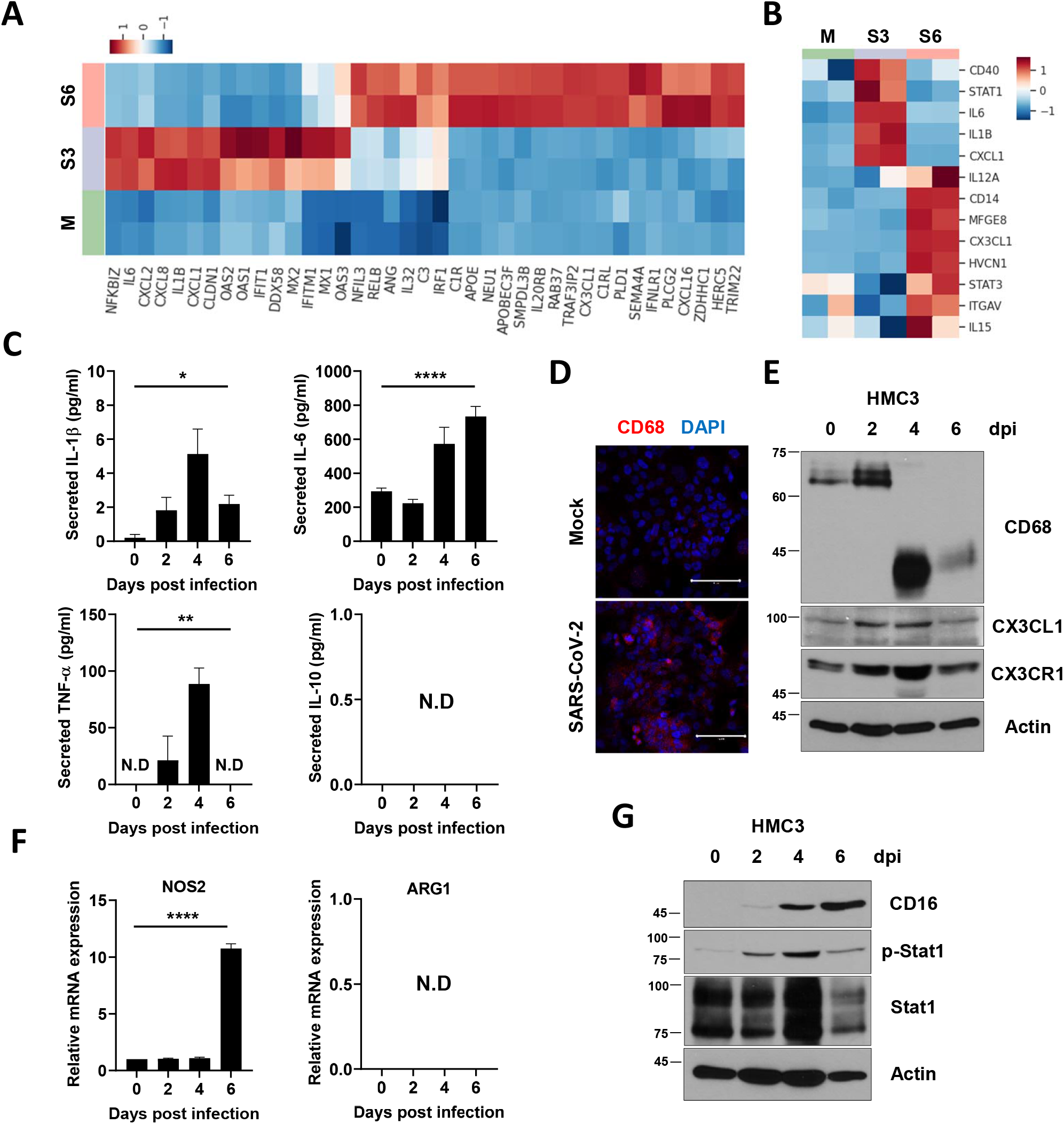
Pro-inflammatory activation and M1 polarization of HMC3 by SARS-CoV-2 infection. **A-B** Heat maps of significantly upregulated genes during SARS-CoV-2 infection enriched in (A) immune response and (B) microglial M1 polarization. **C** The graphs show the measurements of secreted pro-inflammatory cytokines, including IL-1β, IL-6, and TNF-α, and anti-inflammatory cytokine IL-10 by enzyme-linked immunosorbent assay (ELISA). **D** Confocal immunofluorescence images of mock or SARS-CoV-2-infected HMC3 with anti-CD68 antibody. **E** Quantitative analysis of microglial activation markers including CD68, CX3CL1, and CX3CR1 using western blotting. Actin served as the loading control. **F** RT-qPCR analysis of the representative M1 (NOS2; left) and M2 markers (Arginase-1; right). **G** Assessment of proteins of M1 markers including CD16, phopho-Stat1, and Stat1 by western blotting. Actin served as the loading control. Statistically significant differences between the groups were determined using one-way analysis of variance (ANOVA); \**P* < 0.05; \*\**P* < 0.01; \*\*\*\**P* < 0.0001. Symbols represent mean ± SEM.

### Intrinsic and extrinsic death-receptor mediated apoptotic cell death by SARS-CoV-2 infection

As illustrated in Fig. 1, SARS-CoV-2 infection leads to cell death in HMC3. A number of viruses such as the Human Immunodeficiency Virus-1 (HIV-1), Hepatitis C Virus (HCV), and Human Papillomavirus (HPV) activate death receptor (DR)-mediated apoptosis in different ways (39). To address the mechanism of cell death caused by SARS-CoV-2, we thought that infected cells would die through apoptosis, which includes three major ways of programmed cell death (PCD), such as necrosis and pyroptosis. Virtually, the expression of several genes associated with apoptotic process in the DEGs were altered towards that of pro-apoptosis (Fig. 4A). According to the western blotting of the proteins related to the apoptotic process, not only extrinsic death-receptor mediated proteins (Fig. 4B), but also those of the intrinsic pathways (Fig. 4C) of apoptosis were elicited in the infected HMC3. Expression of death receptors such as Fas, DR4, DR5, and TNF receptor 2 (TNFR2), which initiate extrinsic apoptosis, were augmented at 4 and 6 dpi. On the other hand, the expression of an apoptosis regulator, Bcl-2, which suppresses apoptosis, was reduced, and those of Bim, Bid, and Bax, which are essential for the activation of BAX-dependent PCD, were increased. Cleavages of caspase-9 in the intrinsic pathway, caspase-8 in the extrinsic pathway, caspase-3, and poly (ADP-ribose) polymerase (PARP) were observed, demonstrating both pathways of apoptosis (Fig. 4D). Death receptor-mediated apoptosis represents an efficient mechanism by which the virus can induce cell death (39), and ER stress induced by viral infection is mainly involved in the intrinsic pathway (34, 35). Elicited apoptosis in the infected HMC3 was confirmed by Annexin V staining (Fig. 4E). To confirm apoptosis, we measured the survived cells at 6 dpi via crystal violet staining following the treatment of potent and selective inhibitors against Caspases. The treatments of Z-DEVD-FMK (Caspase-3 inhibitor), Z-IETD-FMK (Caspase-8 inhibitor), and Z-VAD-FMK (pan-Caspase inhibitor) significantly blocked the apoptotic cell death (Fig. 4F).

**Figure 4.**
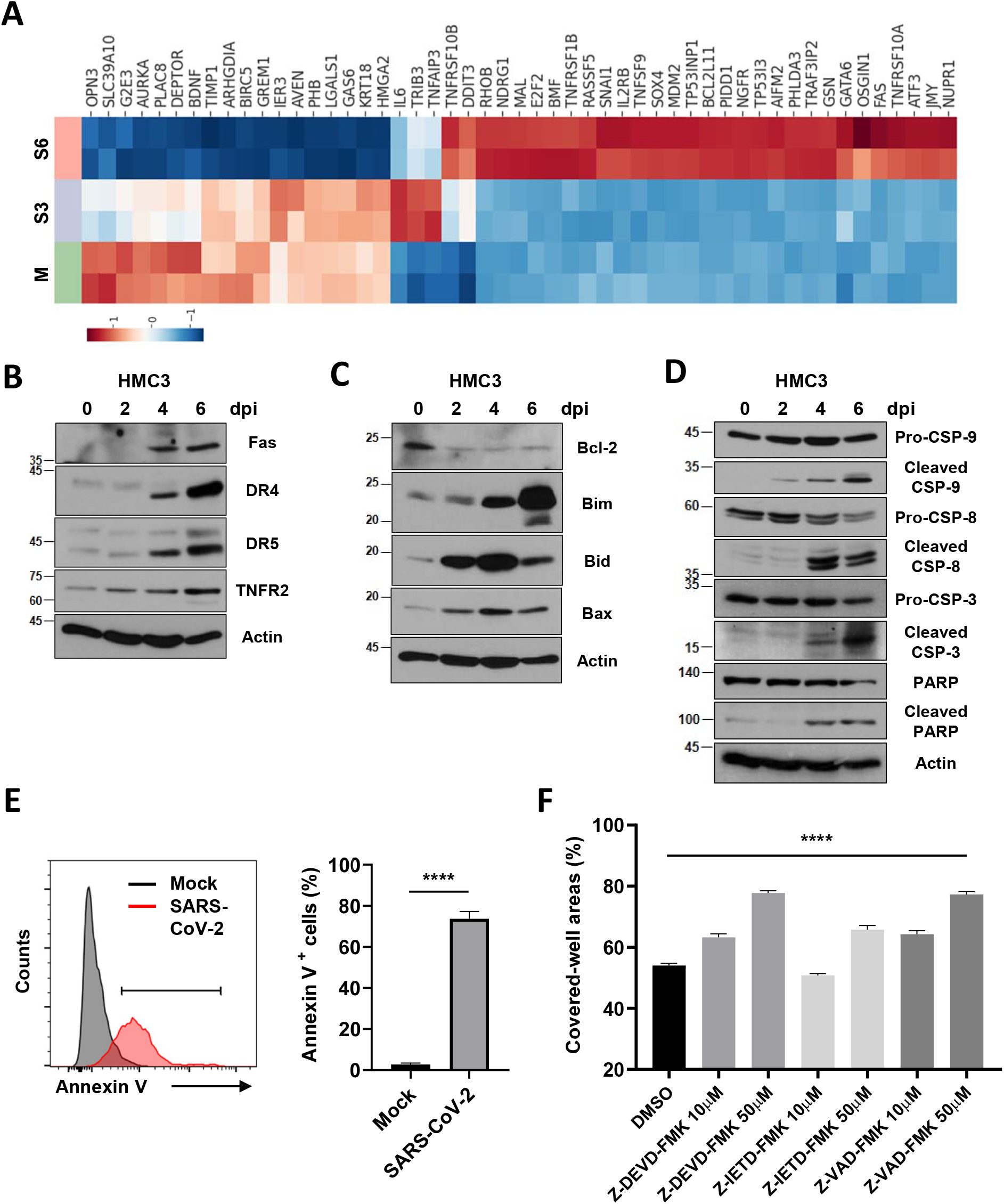
Intrinsic and death receptor (DR)-mediated extrinsic apoptosis in SARS-CoV-2-infected HMC3. **A** A heat map of significantly up- and down-regulated genes during SARS-CoV-2 infection enriched in apoptotic process. **B-D** Extrinsic apoptosis related proteins including Fas, Death receptors (DRs), and tumour necrosis factor receptor 2 (TNFR2) (B); intrinsic apoptosis-associated proteins such as Bcl-2, Bim, Bid, and Bax (C); and caspases and poly (ADP-ribose) polymerase (PARP) (D), which are downstream of both intrinsic and extrinsic apoptosis, were quantitatively analysed using western blotting. Actin served as the loading control. **E** At 3 d post-infection (dpi), the cell surface of the mock or SARS-CoV-2-infected HMC3 was bound with recombinant human Annexin V to detect cells which are in progress of apoptosis by flow cytometry analysis. The histogram peaks indicate mock (gray) and infected (red) cells (Left). The percentage of Annexin V positive cells is shown in the bar graph (Right). **F** The infected HMC3 cells were treated with Z-DEVD-FMK (Caspase-3 inhibitor), Z-IETD-FMK (Caspase-8 inhibitor), and Z-VAD-FMK (pan-Caspase inhibitor) for 6 days at the indicated concentrations. The survived cells were stained with crystal violet and then the percent measurements of the stained cell covered areas was obtained using an ImmunoSpot reader. Statistically significant differences between the groups were determined by Student’s t-test (E) or one-way ANOVA (F); \*\*\*\**P* < 0.0001. Symbols represent means ± SEM.

Since RNA viruses such as Vesicular Stomatitis Virus (VSV), Encephalomyocarditis Virus (EMCV), and Zika virus (ZIKV) promote inflammatory cell death, such as pyroptosis, in most immune cells (40, 41), we assessed the possibility of pyroptosis induced by SARS-CoV-2 infection in HMC3. We observed no changes in the protein expression levels of NLR family pyrin domain containing 3 (NLRP3), or the cleavage of gasdermin D (GSDMD) and caspase-1, which suggests that pyroptosis might not have been promoted in SARS-CoV-2-infected HMC3 (Fig. 5A). The treatments of Ac-FLTD-CMK (Caspase-1 inhibitor) and Belnacasan (VX-765, Caspase-1 inhibitor) as described above also did not recover the cell viability (Fig. 5B). Taken together, these results suggest that SARS-CoV-2 induces cell death in HMC3 through both intrinsic and extrinsic death receptor-mediated apoptosis pathways.

**Figure 5.**
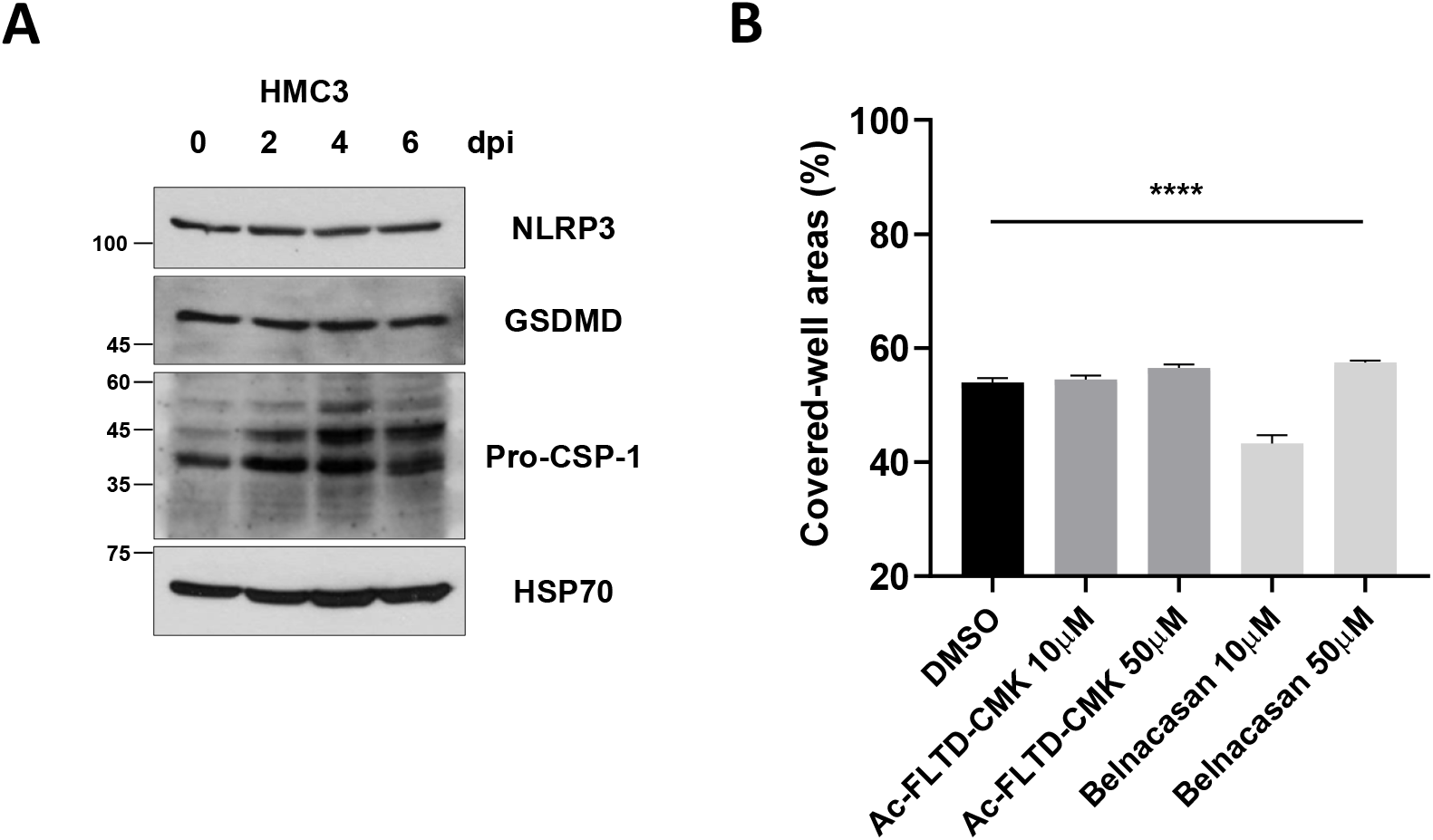
Pyroptosis might not be promoted by SARS-CoV-2 infection in HMC3. **A** After SARS-CoV-2 infection in HMC3, NLRP3, GSDMD, and Caspase-1 proteins were analysed by western blotting. Hsp70 served as a loading control. **B** The SARS-CoV-2-infected HMC3 cells were treated with Caspase-1 inhibitors (Ac-FLTD-CMK and Belnacasan) for 6 days as the indicated concentrations. The survived cells were stained by crystal violet, and then the percent measurements of the stained cell covered areas by ImmunoSpot reader. Statistically significant differences between the groups were determined by one-way ANOVA; \*\*\*\**P* < 0.0001. Symbols represent means ± SEM.

### SARS-CoV-2 can infect microglia of K18-hACE2 mice leading to microgliosis and cell death

To substantiate these viral infection and microglial activation *in vivo* model, we used transgenic mice expressing the human ACE2, driven by the cytokeratin-18 gene promoter (K18-hACE2 mice), which were established for a model of SARS-CoV-2 infection and viral spread through the olfactory pathway (42-44). We inoculated 8-week-old heterozygous male K18-hACE2 mice via the intranasal route with 2 × 10^4^ plaque-forming units (pfu) of SARS-CoV-2. At 6 days post-infection (dpi), infected mice showed a marked weight loss that was approximately 20 % of their body weight (Fig. 6A). The viral RNA was detected from the brains of SARS-CoV-2 infected mice (Fig. 6B). Notably, the co-localization of SARS-CoV-2 spike protein (S) and Iba1, most commonly used marker of microglia, was detected mostly at 6 dpi in the brains (Fig. 6C). From these results, we suggest SARS-CoV-2 infection of microglia *in vivo* model.

**Figure 6.**
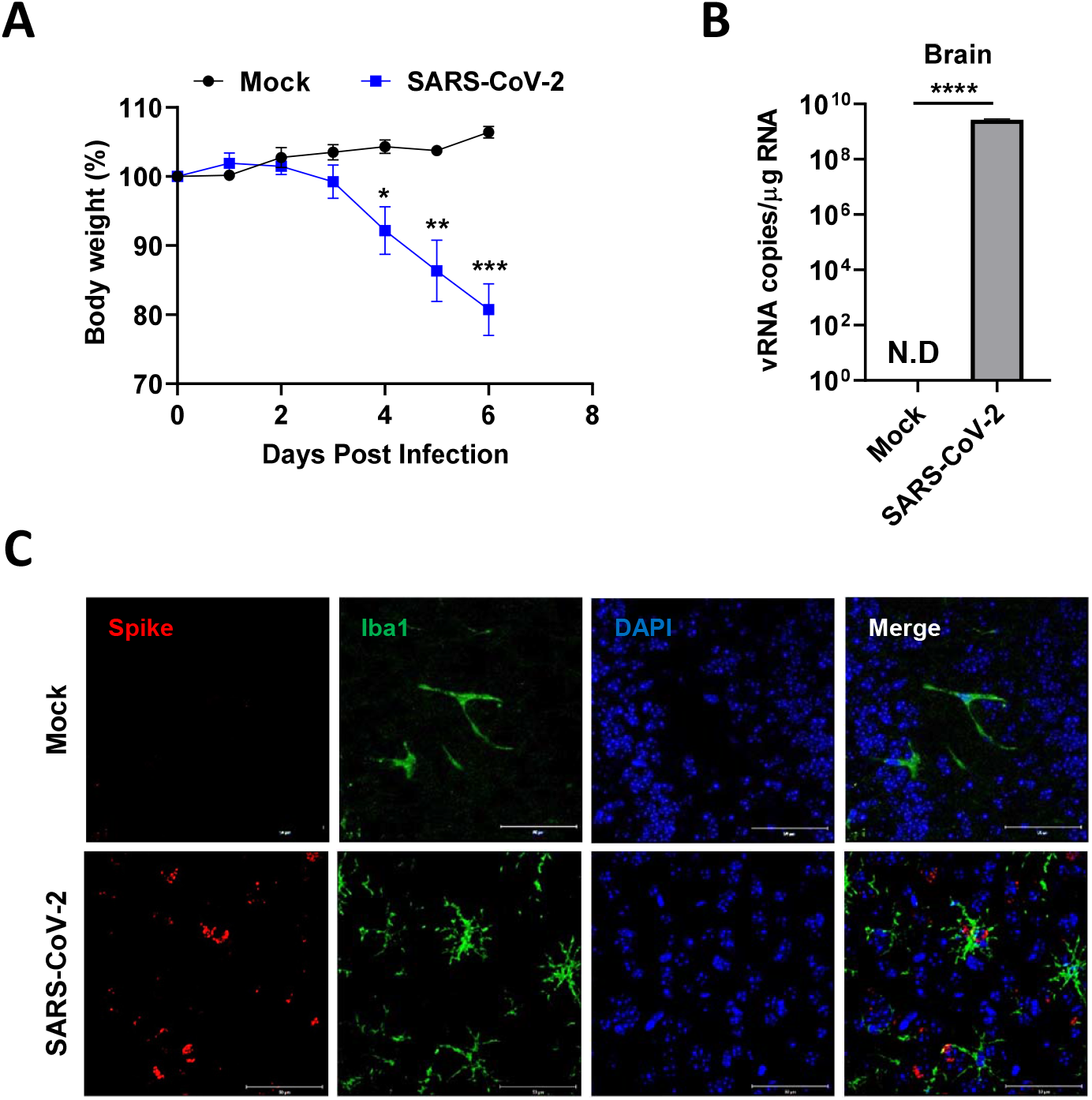
Microglia of K18-hACE2 mice were infected by intranasally administered SARS-CoV-2. **A** The SARS-CoV-2 inoculum (50 μL, 100 MLD50) was intranasally administered to susceptible mouse model (K18-hACE2, n = 4). Their body weight was measured every day (Mock: black; Infected: blue). **B** At 6 d post-infection (dpi), the brain homogenates of the mock or infected mice are used to detect the viral RNA by quantitative real-time polymerase chain reaction (RT-qPCR), and the graph indicates viral RNA copies per microgram of total RNA. **C** The co-localization of SARS-Cov-2 spike protein and microglial Iba1 at 6 dpi in the infected mice by immunofluorescence staining. Scale bars = 50 μm. Statistically significant differences between the groups were determined by multiple student’s *t*-test (A) and Student’s t-test (B); \**P* < 0.05; \*\**P* < 0.01; \*\*\**P* < 0.001; \*\*\*\**P* < 0.0001. symbols represent mean ± SEM.

To confirm that SARS-CoV-2 infection of microglia induces pro-inflammatory activation following cell death in these mice, their brain homogenates were used to isolate leukocytes containing microglia by 30 % and 70 % Percoll discontinuous gradient centrifugation, followed by flow cytometry analysis with cell surface markers of microglia, CD11b and CD45 as illustrated in Fig. 7A. The isolate consisted of three separated populations, including that of lymphocytes (CD11b^-^, CD45^High^), macrophages (CD11b^+^, CD45^High^), and microglia (CD11b^+^, CD45^Low^) (45). While microglia account for most CNS leukocytes of uninfected mice, SARS-CoV-2 infection altered their population (Fig. 7B). Microglia were significantly depopulated by SARS-CoV-2 infection (Fig. 7C). On the other hand, the numbers of infiltrated lymphocytes and macrophages were dramatically increased about seven times (Fig. 7D, 7E). We also found that most microglia were activated with the high expressions of TNF-α and IL-6, compared to the resting microglia of uninfected mice (Fig. 7F-7H). These data indicate that SARS-CoV-2 infection of microglia induces neuroinflammatory processes such as microgliosis, immune cell infiltration, and cell death *in vivo*.

**Figure 7.**
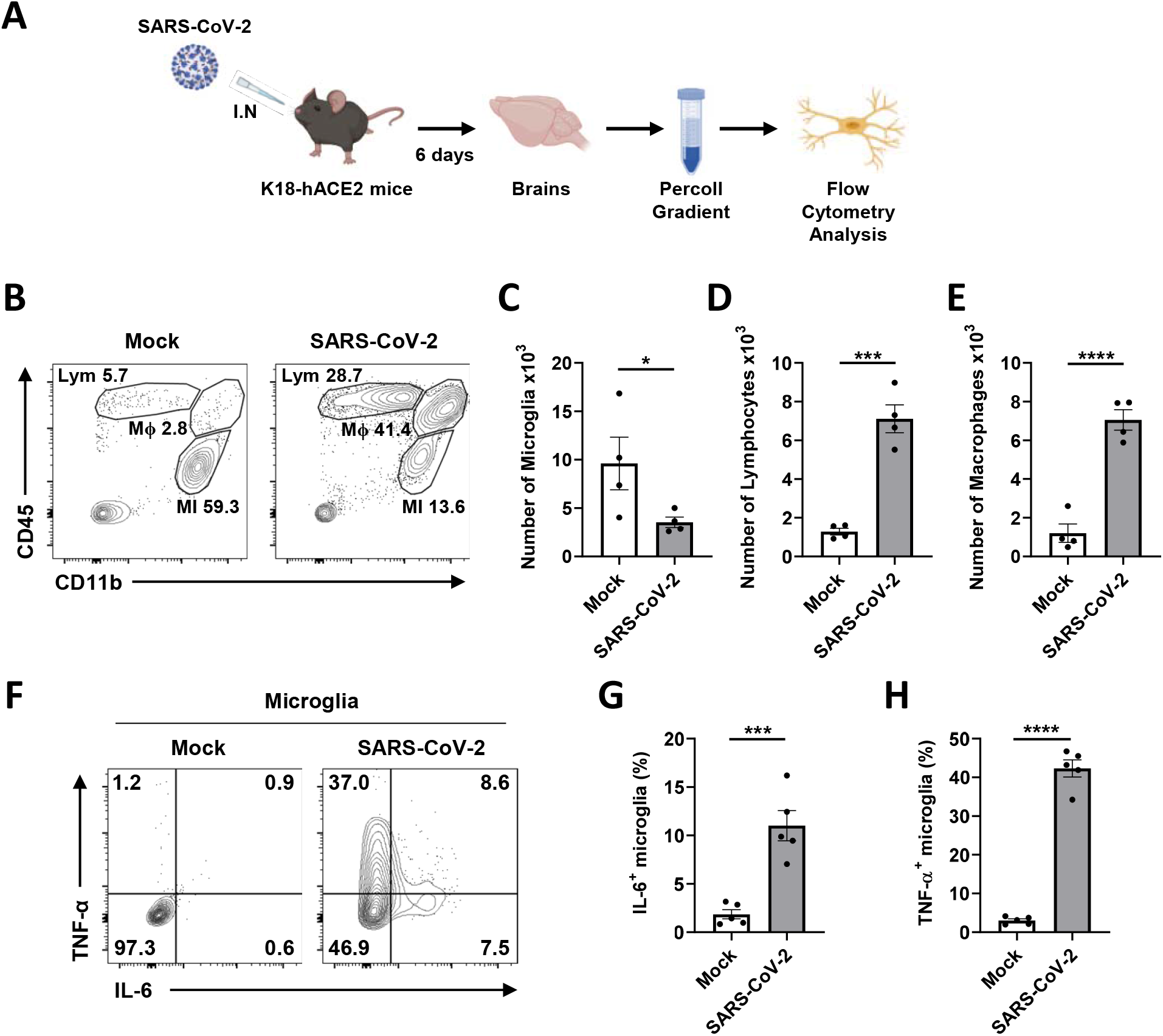
Microglial pro-inflammatory activation and depopulation by SARS-CoV-2 infection in K18-hACE2 mice. **A** Schematic of the experiment for B to H, created with BioRender.com. After six days, brains of mock or SARS-CoV-2-infected mice were extracted and used for Percoll gradient centrifugation to isolate mononuclear cells containing microglia for the flow cytometry analysis. The cellular surface of isolated mononuclear cells was stained with CD11b and CD45 antibodies. **B** Representative flow plot gated on leukocytes shows gating for microglia (MI, CD11b^+^, CD45^Low^), macrophages (Mϕ, CD11b^+^, CD45^High^), and lymphocytes (Lym, CD11b^-^, CD45^High^). **C-E** Bar graphs show the number of microglia (C), lymphocytes (D), and macrophages (E) isolated per brain at 6 dpi. **F** Representative flow plot gated on microglia shows activated microglia with highly expressed IL-6 and TNF-α to separate activated from ramified microglia. **G-H** Bar graphs indicate the percentage of activated microglia, highly expressing IL-6 (G) and TNF-α (H). Statistically significant differences between the groups were determined using Student’s *t-*test; \**P* < 0.05; \*\*\**P* < 0.001; \*\*\*\**P* < 0.0001. Symbols represent means ± standard error of the mean (SEM).

## DISCUSSION

Although COVID-19 is primarily known as a respiratory disease, symptoms related to changes in the peripheral nervous system have recently been reported in the patients (15). There is evidence of viral invasion into the brain through the olfactory or vagal nerve and/or the oral and ophthalmic routes, with trans-synaptic neuronal spread to other brain regions in COVID-19 patients’ autopsy and SARS-CoV-2-infected mice model (25, 29, 46). Microglia, expressing SARS-CoV-2 entry factors, including ACE2 and TMPRSS2 (30), can be infected by the invading SARS-CoV-2 and, subsequently, activated (47). Recently, emerging data have shown the presence of NP expression of Iba1-positive cells in brains of SARS-CoV-2-infected hamsters and even COVID-19 patients with microgliosis (32, 48, 49). Activation of microglia could render individuals more at risk of developing neurological and psychiatric complications, even after complete clinical recovery from the infection (50). Also, dysfunctional or aberrant microglial functions could severely impair cognitive functions, including judgment, decision making, learning, and memory (51). Hence, pro-inflammatory activation by SARS-CoV-2 infection of microglia might have critical outcomes on the short-, moderate-, or long-term neurological and psychiatric consequences of SARS-CoV-2 infection.

In this study, first, we found that SARS-CoV-2 directly infects a human microglial cell line. Regarding the RNA-seq analysis, it was observed that the viral infection induced ER stress and immune responses in the early phase and apoptosis in the late phase. Practically, the microglia, infected with SARS-CoV-2, were activated and polarized toward the M1 phenotype, the mediator of pro-inflammatory responses. In addition, SARS-CoV-2 infection triggered apoptosis as one of the CPE in human microglia through both the intrinsic and extrinsic pathways, further provoking cell death. Indeed, SARS-CoV-2 has been reported to induce both the intrinsic and extrinsic apoptosis in lung tissues and cell lines (52). More recently, SARS-CoV-2-encoded membrane glycoprotein (M) and NP have been demonstrated to trigger apoptosis (53), suggesting a mechanism of apoptosis in SARS-CoV-2-infected microglia. Moreover, we observed SARS-CoV-2 infection of microglia in K18-hACE2 mice, followed by an increase in the number of activated microglia and immune cell accumulation in the brain, and a decrease of the total number of microglia.

Given the co-localization of viral proteins and Iba1 in brains of COVID-19 patients by post-mortem studies (32, 49), we herein suggest a possibility that SARS-CoV-2 directly infects human microglia, inducing the CPE, which is cell death. Also, human microglia could be one of models to study viral pathogenesis. Microglia are important not only for the innate, but also for the adaptive immune responses to pathogen infection of the brain (54). Depletion of microglia can lead to ineffective T cell responses by reduction of the total number and percentage of CD4^+^ and regulatory T cells. Consequently, the depletion of microglia causes increased viral replication and increased neurological manifestations in the brain (55, 56). Therefore, microglial cell death by SARS-CoV-2 infection might be linked to the lack of immune response and consequent increased viral replication, leading to increased neurological manifestations.

The activation of microglia and complement-mediated pathways, which leads to the synthesis of inflammation mediators, is the key element of main inflammatory neurological diseases (57). Virtually, many viruses, including the HIV-1, Herpes Simplex Virus (HSV), and ZIKV, infect microglia and cause neuroinflammation via microglial activation. C3 and its cleavage products attract microglia to gather around the neurons to exert phagocytosis activity and to clear the presynaptic ends (58). The increase in the number of activated microglia via viral infection can have detrimental effects, indirectly by activating astrocytes (59) and T lymphocytes (60, 61) and directly by inducing neuronal damage and death, further contributing to neuronal degeneration (54, 62). In addition, the cytokine storm produced by microglia can lead to increased blood-brain barrier (BBB) breakdown and might be responsible for several neurological symptoms of COVID-19 (47). Thus, microglia could be a potential target for SARS-CoV-2 and could help the spread of the virus in the CNS. In future study, to gain more insights into the SARS-COV-2 neuropathology in COVID-19 patients, it will be important to consider of the relationship between SARS-CoV-2 and microglia. Taken together, our findings indicate that microglia are potential mediators of neurological diseases by SARS-CoV-2 and, consequently, can be targets of therapeutic strategies against neurological disorders in COVID-19 patients.

## MATERIALS AND METHODS

### Cells and virus

Human microglial clone 3 (HMC3) (CRL-3304), Caco-2 (HTB-37), and Vero E6 (CRL-1586) cell lines were purchased from ATCC (Manassas, VA, USA). These cells were maintained in Eagle’s Minimum Essential Medium (EMEM; Welgene, Gyeongsangbuk-do, South Korea) containing 10 % foetal bovine serum (FBS) (Gibco, Waltham, MA, USA) and 1 % Pen/Strep (Gibco). The SARS-CoV-2 Korean strain (GISAID Accession ID: EPI_ISL_407193), isolated from a patient in South Korea, was obtained from Korea Centres for Disease Control and Prevention (KCDC) and propagated in Vero cells (CCL-81, ATCC).

Cells (1 × 10^5^ cells per well) were plated into six-well plates and inoculated with one multiplicity of infection (MOI) SARS-CoV-2 in EMEM containing 2 % FBS on the next day. After incubation for 1 h, the inoculum was removed, and the cells were washed two times. The medium was changed to EMEM containing 10 % FBS. At 2, 4, or 6 dpi, the total cellular RNA was extracted using the RNeasy mini kit (QIAGEN, Hilden, Germany). For the virus neutralization, the inoculum was further incubated with CR3022 neutralizing antibody (ab273073, Abcam, Cambridge, UK) for 1 h. For crystal violet staining, at 6 dpi, cells were stained with 0.5 % crystal violet staining solution in 25 % methanol for 30 min. Then, plates were washed three times with water and dried. The cell-covered areas were measured by the ImmunoSpot reader (CTL, Shaker Heights, OH, USA). The Caspases inhibitors used in this study, including Z-DEVD-FMK (S7312), Z-VAD-FMK (S7023), Z-IETD-FMK (S7314), and Belnacasan (VX-765, S2228), were purchased from Selleckchem (Houston, TX, USA).

### Biosafety

All procedures were performed in a biosafety level 3 (BSL3) or animal BSL3 facility for SARS-CoV-2-related experiments, with approval from the Korea Research Institute of Chemical Technology (KRICT) and by personnel equipped with powered air-purifying respirators.

### Mice

Eight-week-old male B6.Cg-Tg(K18-hACE2)2Prlmn/J mice were purchased from the Jackson Laboratory and maintained in a biosafety level 2 (BSL-2) animal facility in the Korea Research Institute of Chemical Technology (KRICT). All protocols were approved by the Institutional Animal Care and Use Committee (Protocol ID 8A-M6, IACUC ID 2021-8A-02-01 & 2021-8A-03-03).

Virus inoculations (SARS-CoV-2, 2 × 10^4^ pfu) were performed by the intranasal route (I.N.) under anaesthesia using isoflurane in a BSL-3 animal facility, and all efforts were made to minimise animal suffering. Body weights were measured everyday post-infection.

### Microglia isolation

Isolation of microglia in mice was performed following the protocols previously described (63, 64), with minimal modifications. Mock or SARS-CoV-2 infected mice were anesthetised by isoflurane, followed by perfusion with 10 or 20 ml of cold 1× DPBS (Gibco) into the left ventricle to remove blood from the tissues. Brains were transferred to a six-well plate containing cold Hanks’ Balanced Salt Solution (HBSS) (Gibco) and the plates were kept on ice. The generation of brain cell suspension by a 70-μm pore sized-cell strainer (SPL, Gyeonggi-do, South Korea) was made in 10 ml per brain of digestion cocktail containing 0.5 mg/ml DNase I (Roche, Basel, Switzerland) and 1 mg/ml Collagenase A (Roche) in HBSS. The suspension was incubated at room temperature for 30 min, followed centrifugation for 7 min at 300 × *g*, 18 °C. The cell pellet was resuspended with 30 % Percoll (Sigma-Aldrich, St. Louis, MO, USA) in HBSS, and then was slowly layered over 70 % Percoll in HBSS in a 15 ml-conical tube. About 2 ml of interphase volume was collected to a new tube after gradient centrifugation for 40 min at 200 × *g*, 18 °C. Isolated mononuclear cells were washed three more times in a volume of 500 μL of HBSS containing 0.01 M HEPES (Gibco), using a micro-centrifuge for 7 min at 600 × *g*, 4 °C.

### Flow cytometry analysis

Isolated brain mononuclear cells from the mock or SARAS-CoV-2-infected mice in cell staining buffer (phosphate-buffered saline [PBS] with 1 % FBS and 0.09 % NaN3) were stained for 30 min with fluorescence-conjugated antibodies, namely, Brilliant Violet 421 anti-mouse/human CD11b Antibody (101236, BioLegend, San Diego, CA, USA), PE/Cyanine7 anti-mouse CD45 Antibody (103114, BioLegend), APC anti-mouse TNF-α Antibody (506307, BioLegend), FITC anti-mouse IL-6 Monoclonal Antibody (MP5-20F3) (11-7061-82, eBioscience, San Diego, CA, USA), FITC anti-human ACE2 Antibody (NBP2-7211F, Novus Biologicals, Centennial, CO, USA), and Alexa 647 anti-Iba1 Antibody (78060S, Cell signalling Technology, Danvers, MA, USA). Cells were then analysed by FACSAria III sorter (BD Biosciences, San Jose, CA, USA), and data was analysed by the FlowJo software (BD Biosciences). All fluorochromes were compensated. The total leukocyte population was gated for microglia (CD11b^+^, CD45^Low^). For Annexin V staining, mock or SARS-CoV-2-infected HMC3 were stained with FITC-recombinant human Annexin V, following the protocols of Annexin V-FITC Apoptosis Detection Kit (BMS500FI-20, eBioscience).

### RT-qPCR

Quantitative RT-PCR (QuantStudio 3, Applied Biosystems, Foster City, CA, USA) was performed with one-step Prime script III RT-qPCR mix (Takara, Japan). The viral RNA of NP was detected by 2019-nCoV-N1 probe (Cat#10006770, Integrated DNA Technologies, Coralville, IA, USA). The IL-1β, IL-6, IL-12, TNF-α, IFN-β, IFN-λ1, NOS2, and Arginae-1 genes were detected by individual customized probes (Integrated DNA Technologies).

### Focus forming assay

The cell culture media serially diluted in EMEM containing 2 % FBS was added to 4 × 10^4^ Vero E6 cells plated on 96-well plates. After incubation for 8 h at 37 °C, cells were washed and fixed with 4 % formaldehyde solution. Cells were stained with the anti-SARS-CoV-2 NP antibody (40143-R001, Sino biological, Beijing, China) and a secondary horseradish peroxidase-conjugated goat anti-rabbit IgG (Bio-Rad, Hercules, CA). The signal was developed using an insoluble tetramethybenzidine (TMB) substrate (Promega, Madison, WI, USA), and the number of infected cells were counted using an ImmunoSpot reader (CTL, Shaker Heights, OH).

### RNA-seq and analysis

Sequencing library was prepared with TruSeq Stranded mRNA Sample Prep Kit and sequenced on NovaSeq 6000 (Illumina, San Diego, CA, USA), yielding more than 6G bases of sequences for each sample. From the sequenced reads, adaptor sequences were removed using Cutadapt (version 3.1) (65) and aligned to the hybrid reference genomes of human (GRCh38.p13_ENS100) and SARS-CoV-2 (ASM985889_v3) with STAR aligner (version 2.7.6a) (66). Aligned reads were quantified in gene level by HTSeq (version 0.13.5) (67) with “intersection-nonempty” mode. Genes with lower than five counts for the total count per gene were removed for further analyses. Differentially expressed gene analysis were processed with DESeq2 (version 1.30.1) (68) using abs (log2 fold change) > 1 and adjusted P-value (Benjamini-Hochberg) < 0.01 as cut-off. Multidimensional scaling analysis were performed with clustermap function in python seaborn package (version 0.11.1) using genes with mean FPKM > 1 among the samples and transformed to log2(FPKM+1). Over-representation analysis of the DEGs enriched to GO Biological Process 2018 with EnrichR (69) with adjusted P-value (Benjamini-Hochberg) < 0.05 cut-off. Network analysis were presented by Metascape (70) using the GO Biological Process gene sets.

### Immunofluorescence assay

After transcardial perfusion with cold 4 % paraformaldehyde in PBS, brain tissues of SARS-CoV-2 infected K18-hACE2 mice were dissected and fixed by immersion in 4 % paraformaldehyde in PBS overnight at 4 °C. The brain sections (30 μm thickness) were permeabilized with 0.2 % Triton X-100 in 1 % BSA/PBS for 30 min, washed in PBS, and blocked with 0.5 % BSA in PBS for 15 min, followed by incubation overnight at 4 °C with primary antibodies, namely, anti-SARS-CoV-2 S (40150-T62-COV2, Sino Biological), and anti-Iba1/AIF1 (MABN92, Merck Millipore, Burlington, MA, USA). After washing twice, further incubation was carried out with Alexa Fluor 488-conjugated anti-rabbit antibody (A32731, Thermo Fisher Scientific, Waltham, MA, USA) and Alexa Fluor 594-conjugated anti-mouse antibody (A32744, Thermo Fisher Scientific). Immunofluorescence was observed by confocal microscopy (LSM700, Carl Zeiss, Oberkochen, Germany).

For the cell lines, the SARS-CoV-2-infected cells were fixed with 4 % paraformaldehyde in PBS overnight at 4 °C, and then permeabilized with 0.5 % Triton X-100 in PBS for 10 min, followed by washing thrice with PBS. Blocking buffer (0.1 % Tween 20, 1 % bovine serum albumin [BSA] in PBS) was added to remove non-specific binding. Cells were immunostained overnight at room temperature with primary antibodies, namely, anti-dsRNA J2 (MABE1134, Sigma-Aldrich), anti-SARS-CoV-2 NP (40143-R019, Sino Biological), anti-SARS-CoV-2 S (40150-T62-COV2, Sino Biological), and anti-CD68 (sc-17832, Santa Cruz Biotechnology, Dallas, TX, USA). After washing thrice, further incubation was carried out with Alexa Fluor 488-conjugated anti-rabbit antibody (A32731, Thermo Fisher Scientific) and Alexa Fluor 594-conjugated anti-mouse antibody (A32744, Thermo Fisher Scientific). Immunofluorescence was observed by the confocal microscopy.

### ELISA

The culture supernatants were collected from infected cells and used for detection of IL-1β, IL-6, IL-10, and TNF-α. Each cytokine was determined by the corresponding ELISA kit (IL-1β, K0331800; IL-6, K0331194; IL-10, K0331123; TNF-α, K0331131; Komabiotech, Seoul, South Korea), following the manufacturer’s instructions.

### Western blotting

Cells were lysed in radioimmunoprecipitation (RIPA) buffer (Thermo Fisher Scientific), and proteins in the lysate were separated in a denaturing polyacrylamide gel and transferred to a polyvinylidene fluoride (PVDF) membrane (Merck Millipore, Burlington, MA, USA). The membrane was incubated with 5 % skim milk (BD Biosciences) in Tris-buffered saline with 0.1 % Tween 20 (TBST) buffer and the primary antibodies, namely, anti-SARS-CoV-2 NP (40143-R001, Sino biological), anti-CD68 (sc-17832, Santa Cruz Biotechnology) anti-GSDMDC1 (sc-81868, Santa Cruz Biotechnology), anti-Actin (sc-47778, Santa Cruz Biotechnology), anti-Hsp70 (sc-24, Santa Cruz Biotechnology), anti-CX3CL1 (ab25088, Abcam), anti-CX3CR1 (ab8021, Abcam), anti-CD16 (80006S, Cell signalling Technology), anti-phospho-Stat1 (9167S, Cell signalling Technology), anti-Stat1 (14994S, Cell signalling Technology), anti-Fas (4233T, Cell signalling Technology), anti-DR4 (42533T, Cell signalling Technology), anti-DR5 (8074T, Cell signalling Technology), anti-TNFR2 (3727T, Cell signalling Technology), anti-Bcl-2 (4223T, Cell signalling Technology), anti-Bim (2933T, Cell signalling Technology), anti-Bid (2002T, Cell signalling Technology), anti-Bax (5023T, Cell signalling Technology), anti-Caspase-9 (9502S, Cell signalling Technology), anti-Caspase-8 (9746S, Cell signalling Technology), anti-Caspase-3 (9665S, Cell signalling Technology), anti-Caspase-1 (3866S, Cell Signalling Technology), anti-NLRP3 (15101S, Cell Signalling Technology), and anti-PARP (9542S, Cell signalling Technology). Horseradish peroxidase (HRP)-conjugated secondary antibodies from Bio-Rad and ECL reagents (Thermo Fisher Scientific) were used for protein detection.

### Statistical analysis

All experiments were performed at least three times. All data were analysed using the GraphPad Prism 8.0 software (GraphPad Software, San Diego, CA, USA). *P* < 0.05 was considered statistically significant. Specific analysis methods are described in the figure legends.

## Acknowledgments

The SARS-CoV-2 (NCCP43326) was kindly provided by the National Culture Collection for Pathogens at the Korea Centers for Disease Control and Prevention. This work was supported by the National Research Foundation of Korea (NRF) grant, funded by the Ministry of Education, Science, and Technology (MIST) of the Korean government (2020R1C1C1003379) and the National Research Council of Science & Technology (NST) grant, funded by the Korean government (MSIP) (CRC-16-01-KRICT).

## Author contributions

Conceptualization: G.U.J., and Y.-C.K.; methodology: G.U.J., K.-D.K., J.K., W.H.S. and Y.-C.K.; investigation: G.U.J., J.R., K.-D.K., Y.C.C., G.Y.Y., S.L., and I.H.; writing: G.U.J. and J.R.; review and editing: G.U.J., K.-D.K., W.H.S., J.K., J-Y.L., and Y-C.K.; funding acquisition: Y.-C.K.; and supervision: Y.-C.K.

## References

1. V’kovski P, Kratzel A, Steiner S, Stalder H, Thiel V. 2021. Coronavirus biology and replication: implications for SARS-CoV-2. Nature Reviews Microbiology 19:155–170.

2. Hu B, Guo H, Zhou P, Shi Z-L. 2020. Characteristics of SARS-CoV-2 and COVID-19. Nature Reviews Microbiology:1–14.

3. Fotuhi M, Mian A, Meysami S, Raji CA. 2020. Neurobiology of COVID-19. Journal of Alzheimer’s disease:1–17.

4. Varatharaj A, Thomas N, Ellul MA, Davies NW, Pollak TA, Tenorio EL, Sultan M, Easton A, Breen G, Zandi M. 2020. Neurological and neuropsychiatric complications of COVID-19 in 153 patients: a UK-wide surveillance study. The Lancet Psychiatry 7:875–882.

5. Politi LS, Salsano E, Grimaldi M. 2020. Magnetic resonance imaging alteration of the brain in a patient with coronavirus disease 2019 (COVID-19) and anosmia. JAMA neurology 77:1028–1029.

6. Reichard RR, Kashani KB, Boire NA, Constantopoulos E, Guo Y, Lucchinetti CF. 2020. Neuropathology of COVID-19: a spectrum of vascular and acute disseminated encephalomyelitis (ADEM)-like pathology. Acta neuropathologica 140:1–6.

7. Gautier J-F, Ravussin Y. 2020. A new symptom of COVID-19: loss of taste and smell. Obesity (Silver Spring) 28:848.

8. Giacomelli A, Pezzati L, Conti F, Bernacchia D, Siano M, Oreni L, Rusconi S, Gervasoni C, Ridolfo AL, Rizzardini G. 2020. Self-reported olfactory and taste disorders in patients with severe acute respiratory coronavirus 2 infection: a cross-sectional study. Clinical Infectious Diseases 71:889–890.

9. Poyiadji N, Shahin G, Noujaim D, Stone M, Patel S, Griffith B. 2020. COVID-19–associated acute hemorrhagic necrotizing encephalopathy: imaging features. Radiology 296:E119–E120.

10. Li Z, Liu T, Yang N, Han D, Mi X, Li Y, Liu K, Vuylsteke A, Xiang H, Guo X. 2020. Neurological manifestations of patients with COVID-19: potential routes of SARS-CoV-2 neuroinvasion from the periphery to the brain. Frontiers of medicine:1–9.

11. Chen T, Wu D, Chen H, Yan W, Yang D, Chen G, Ma K, Xu D, Yu H, Wang H. 2020. Clinical characteristics of 113 deceased patients with coronavirus disease 2019: retrospective study. bmj 368.

12. Phua J, Weng L, Ling L, Egi M, Lim C-M, Divatia JV, Shrestha BR, Arabi YM, Ng J, Gomersall CD. 2020. Intensive care management of coronavirus disease 2019 (COVID-19): challenges and recommendations. The Lancet Respiratory Medicine 8:506–517.

13. Helms J, Kremer S, Merdji H, Clere-Jehl R, Schenck M, Kummerlen C, Collange O, Boulay C, Fafi-Kremer S, Ohana M. 2020. Neurologic features in severe SARS-CoV-2 infection. New England Journal of Medicine 382:2268–2270.

14. Oxley TJ, Mocco J, Majidi S, Kellner CP, Shoirah H, Singh IP, De Leacy RA, Shigematsu T, Ladner TR, Yaeger KA. 2020. Large-vessel stroke as a presenting feature of Covid-19 in the young. New England Journal of Medicine 382:e60.

15. Mao L, Jin H, Wang M, Hu Y, Chen S, He Q, Chang J, Hong C, Zhou Y, Wang D. 2020. Neurologic manifestations of hospitalized patients with coronavirus disease 2019 in Wuhan, China. JAMA neurology 77:683–690.

16. Block ML, Zecca L, Hong J-S. 2007. Microglia-mediated neurotoxicity: uncovering the molecular mechanisms. Nature Reviews Neuroscience 8:57–69.

17. Fetler L, Amigorena S. 2005. Brain under surveillance: the microglia patrol. Science 309:392–393.

18. Sochocka M, Diniz BS, Leszek J. 2017. Inflammatory response in the CNS: friend or foe? Molecular neurobiology 54:8071–8089.

19. Cho BP, Song DY, Sugama S, Shin DH, Shimizu Y, Kim SS, Kim YS, Joh TH. 2006. Pathological dynamics of activated microglia following medial forebrain bundle transection. Glia 53:92–102.

20. Brown GC, Vilalta A. 2015. How microglia kill neurons. Brain research 1628:288–297.

21. Polazzi E, Contestabile A. 2002. Reciprocal interactions between microglia and neurons: from survival to neuropathology. Reviews in the Neurosciences 13:221–242.

22. Zecca L, Zucca F, Albertini A, Rizzio E, Fariello R. 2006. A proposed dual role of neuromelanin in the pathogenesis of Parkinson’s disease. Neurology 67:S8–S11.

23. McGeer PL, Rogers J, McGeer EG. 2006. Inflammation, anti-inflammatory agents and Alzheimer disease: the last 12 years. Journal of Alzheimer’s Disease 9:271–276.

24. Kim YS, Joh TH. 2006. Microglia, major player in the brain inflammation: their roles in the pathogenesis of Parkinson’s disease. Experimental & molecular medicine 38:333–347.

25. Matschke J, Lütgehetmann M, Hagel C, Sperhake JP, Schröder AS, Edler C, Mushumba H, Fitzek A, Allweiss L, Dandri M. 2020. Neuropathology of patients with COVID-19 in Germany: a post-mortem case series. The Lancet Neurology 19:919–929.

26. Schurink B, Roos E, Radonic T, Barbe E, Bouman CS, de Boer HH, de Bree GJ, Bulle EB, Aronica EM, Florquin S. 2020. Viral presence and immunopathology in patients with lethal COVID-19: a prospective autopsy cohort study. The Lancet Microbe 1:e290–e299.

27. Al-Dalahmah O, Thakur KT, Nordvig AS, Prust ML, Roth W, Lignelli A, Uhlemann A-C, Miller EH, Kunnath-Velayudhan S, Del Portillo A. 2020. Neuronophagia and microglial nodules in a SARS-CoV-2 patient with cerebellar hemorrhage. Acta Neuropathologica Communications 8:1–7.

28. Song E, Zhang C, Israelow B, Lu-Culligan A, Prado AV, Skriabine S, Lu P, Weizman O-E, Liu F, Dai Y. 2021. Neuroinvasion of SARS-CoV-2 in human and mouse brain. Journal of Experimental Medicine 218:e20202135.

29. Meinhardt J, Radke J, Dittmayer C, Franz J, Thomas C, Mothes R, Laue M, Schneider J, Brünink S, Greuel S. 2021. Olfactory transmucosal SARS-CoV-2 invasion as a port of central nervous system entry in individuals with COVID-19. Nature neuroscience 24:168–175.

30. Singh M, Bansal V, Feschotte C. 2020. A single-cell RNA expression map of human coronavirus entry factors. Cell reports 32:108175.

31. Bengsch B, Schwabenland M, Salié H, Tanevski J, Killmer S, Matschke J, Püschel K, Mei H, Boettler T, Neumann-Haefelin C. 2020. Deep spatial profiling of COVID19 brains reveals neuroinflammation by compartmentalized local immune cell interactions and targets for intervention.

32. Cama VF, Marín-Prida J, Acosta-Rivero N, Acosta EF, Díaz LO, Casadesús AV, Fernández-Marrero B, Gilva-Rodríguez N, Cremata-García D, Cervantes-Llanos M. 2021. The microglial NLRP3 inflammasome is involved in human SARS-CoV-2 cerebral pathogenicity: A report of three post-mortem cases. Journal of Neuroimmunology:577728.

33. Wurtz N, Penant G, Jardot P, Duclos N, La Scola B. 2021. Culture of SARS-CoV-2 in a panel of laboratory cell lines, permissivity, and differences in growth profile. European Journal of Clinical Microbiology & Infectious Diseases 40:477–484.

34. Jheng J-R, Ho J-Y, Horng J-T. 2014. ER stress, autophagy, and RNA viruses. Frontiers in microbiology 5:388.

35. He B. 2006. Viruses, endoplasmic reticulum stress, and interferon responses. Cell Death & Differentiation 13:393–403.

36. Szegezdi E, Logue SE, Gorman AM, Samali A. 2006. Mediators of endoplasmic reticulum stress-induced apoptosis. EMBO reports 7:880–885.

37. Jiang CT, Wu WF, Deng YH, Ge JW. 2020. Modulators of microglia activation and polarization in ischemic stroke. Molecular medicine reports 21:2006–2018.

38. Hopperton K, Mohammad D, Trépanier M, Giuliano V, Bazinet R. 2018. Markers of microglia in post-mortem brain samples from patients with Alzheimer’s disease: a systematic review. Molecular psychiatry 23:177–198.

39. Zhou X, Jiang W, Liu Z, Liu S, Liang X. 2017. Virus infection and death receptor-mediated apoptosis. Viruses 9:316.

40. da Costa LS, Outlioua A, Anginot A, Akarid K, Arnoult D. 2019. RNA viruses promote activation of the NLRP3 inflammasome through cytopathogenic effect-induced potassium efflux. Cell death & disease 10:1–15.

41. Wang W, Li G, Wu D, Luo Z, Pan P, Tian M, Wang Y, Xiao F, Li A, Wu K. 2018. Zika virus infection induces host inflammatory responses by facilitating NLRP3 inflammasome assembly and interleukin-1β secretion. Nature communications 9:1–16.

42. McCray PB, Pewe L, Wohlford-Lenane C, Hickey M, Manzel L, Shi L, Netland J, Jia HP, Halabi C, Sigmund CD. 2007. Lethal infection of K18-hACE2 mice infected with severe acute respiratory syndrome coronavirus. Journal of virology 81:813–821.

43. Netland J, Meyerholz DK, Moore S, Cassell M, Perlman S. 2008. Severe acute respiratory syndrome coronavirus infection causes neuronal death in the absence of encephalitis in mice transgenic for human ACE2. Journal of virology 82:7264–7275.

44. Winkler ES, Bailey AL, Kafai NM, Nair S, McCune BT, Yu J, Fox JM, Chen RE, Earnest JT, Keeler SP. 2020. SARS-CoV-2 infection of human ACE2-transgenic mice causes severe lung inflammation and impaired function. Nature immunology 21:1327–1335.

45. Rangaraju S, Raza SA, Li NXA, Betarbet R, Dammer EB, Duong D, Lah JJ, Seyfried NT, Levey AI. 2018. Differential phagocytic properties of CD45low microglia and CD45high brain mononuclear phagocytes—activation and age-related effects. Frontiers in immunology 9:405.

46. Cantuti-Castelvetri L, Ojha R, Pedro LD, Djannatian M, Franz J, Kuivanen S, van der Meer F, Kallio K, Kaya T, Anastasina M. 2020. Neuropilin-1 facilitates SARS-CoV-2 cell entry and infectivity. Science 370:856–860.

47. Vargas G, Geraldo LHM, Salomão N, Paes MV, Lima FRS, Gomes FCA. 2020. Severe acute respiratory syndrome coronavirus 2 (SARS-CoV-2) and glial cells: Insights and perspectives. Brain, behavior, & immunity-health:100127.

48. de Melo GD, Lazarini F, Levallois S, Hautefort C, Michel V, Larrous F, Verillaud B, Aparicio C, Wagner S, Gheusi G. 2021. COVID-19–related anosmia is associated with viral persistence and inflammation in human olfactory epithelium and brain infection in hamsters. Science Translational Medicine 13.

49. Schwabenland M, Salié H, Tanevski J, Killmer S, Lago MS, Schlaak AE, Mayer L, Matschke J, Püschel K, Fitzek A. 2021. Deep spatial profiling of human COVID-19 brains reveals neuroinflammation with distinct microanatomical microglia-T-cell interactions. Immunity 54:1594–1610. e11.

50. Tremblay M-È. 2020. A Diversity of Cell Types, Subtypes and Phenotypes in the Central Nervous System: The Importance of Studying Their Complex Relationships. Frontiers in Cellular Neuroscience 14.

51. Tay TL, Béchade C, D’Andrea I, St-Pierre M-K, Henry MS, Roumier A, Tremblay M-E. 2018. Microglia gone rogue: impacts on psychiatric disorders across the lifespan. Frontiers in molecular neuroscience 10:421.

52. Liu Y, Garron TM, Chang Q, Su Z, Zhou C, Qiu Y, Gong EC, Zheng J, Yin YW, Ksiazek T. 2021. Cell-type apoptosis in lung during SARS-CoV-2 infection. Pathogens 10:509.

53. Ren Y, Wang A, Fang Y, Shu T, Wu D, Wang C, Huang M, Min J, Jin L, Zhou W. 2021. SARS-CoV-2 membrane glycoprotein M triggers apoptosis with the assistance of nucleocapsid protein N in cells. Frontiers in cellular and infection microbiology:627.

54. Klein RS, Garber C, Funk KE, Salimi H, Soung A, Kanmogne M, Manivasagam S, Agner S, Cain M. 2019. Neuroinflammation during RNA viral infections. Annual review of immunology 37:73–95.

55. Wheeler DL, Sariol A, Meyerholz DK, Perlman S. 2018. Microglia are required for protection against lethal coronavirus encephalitis in mice. The Journal of clinical investigation 128:931–943.

56. Mangale V, Syage AR, Ekiz HA, Skinner DD, Cheng Y, Stone CL, Brown RM, O’Connell RM, Green KN, Lane TE. 2020. Microglia influence host defense, disease, and repair following murine coronavirus infection of the central nervous system. Glia 68:2345–2360.

57. Degan D, Ornello R, Tiseo C, Carolei A, Sacco S, Pistoia F. 2018. The role of inflammation in neurological disorders. Current pharmaceutical design 24:1485–1501.

58. Chen Z, Zhong D, Li G. 2019. The role of microglia in viral encephalitis: a review. Journal of neuroinflammation 16:1–12.

59. Tremblay M-È, Stevens B, Sierra A, Wake H, Bessis A, Nimmerjahn A. 2011. The role of microglia in the healthy brain. Journal of Neuroscience 31:16064–16069.

60. Zimmermann J, Hafezi W, Dockhorn A, Lorentzen EU, Krauthausen M, Getts DR, Müller M, Kühn JE, King NJ. 2017. Enhanced viral clearance and reduced leukocyte infiltration in experimental herpes encephalitis after intranasal infection of CXCR3-deficient mice. Journal of neurovirology 23:394–403.

61. Garber C, Soung A, Vollmer LL, Kanmogne M, Last A, Brown J, Klein RS. 2019. T cells promote microglia-mediated synaptic elimination and cognitive dysfunction during recovery from neuropathogenic flaviviruses. Nature neuroscience 22:1276–1288.

62. Trzeciak A, Lerman YV, Kim T-H, Kim MR, Mai N, Halterman MW, Kim M. 2019. Long-term microgliosis driven by acute systemic inflammation. The Journal of Immunology 203:2979–2989.

63. Cardona AE, Huang D, Sasse ME, Ransohoff RM. 2006. Isolation of murine microglial cells for RNA analysis or flow cytometry. Nature protocols 1:1947–1951.

64. Garcia JA, Cardona SM, Cardona AE. 2014. Isolation and analysis of mouse microglial cells. Current protocols in immunology 104:14.35. 1-14.35. 15.

65. Martin M. 2011. Cutadapt removes adapter sequences from high-throughput sequencing reads. EMBnet journal 17:10–12.

66. Dobin A, Davis CA, Schlesinger F, Drenkow J, Zaleski C, Jha S, Batut P, Chaisson M, Gingeras TR. 2013. STAR: ultrafast universal RNA-seq aligner. Bioinformatics 29:15–21.

67. Anders S, Pyl PT, Huber W. 2015. HTSeq—a Python framework to work with high-throughput sequencing data. Bioinformatics 31:166–169.

68. Love MI, Huber W, Anders S. 2014. Moderated estimation of fold change and dispersion for RNA-seq data with DESeq2. Genome biology 15:1–21.

69. Kuleshov MV, Jones MR, Rouillard AD, Fernandez NF, Duan Q, Wang Z, Koplev S, Jenkins SL, Jagodnik KM, Lachmann A. 2016. Enrichr: a comprehensive gene set enrichment analysis web server 2016 update. Nucleic acids research 44:W90–W97.

70. Zhou Y, Zhou B, Pache L, Chang M, Khodabakhshi AH, Tanaseichuk O, Benner C, Chanda SK. 2019. Metascape provides a biologist-oriented resource for the analysis of systems-level datasets. Nature communications 10:1–10.

